# Identification of Essential Temperature-Stressed Genes From *Apis mellifera* Hypopharyngeal Glands Transcriptomes Under Variable Temperatures

**DOI:** 10.1101/2023.11.22.568201

**Authors:** Abdulkadir Yusif Maigoro, Jeong Hyeon Lee, Sujin Lee, Hyung Wook Kwon

## Abstract

Temperature is one of the essential abiotic factors required for honey bee survival and pollination. It affects honey bee physiology, behavior, and expression of related genes. Also, considered one of the major factors contributing to colony collapse disease (CCD). In this research, RNA-seq analysis was performed using hypopharyngeal glands (HGs) tissue at low (18 °C), high (25 °C), and regular (22 °C) temperatures. Differentially expressed genes (DEGs) were identified after comparing the three groups with one another based on temperature differences using DESeq analysis. 5196 common DEGs (cDEGs) were identified among the groups. They are highly enriched in RNA processing and RNA metabolism process while the KEGG pathway enrichment analysis showed that the cDEGs are enriched in longevity regulating pathway, MAPK signaling pathway-fly, and Glycerophospholipid metabolism. Further, 360 temperature-stressed genes identified are highly enriched in translation, oxidative activity, and ribonucleoprotein complex. The enriched KEGG pathway includes ribosome, oxidative phosphorylation, fatty acid metabolism, and citrate cycle (TCA cycle). All the top ten (10) hub genes among the 360 temperature-stressed genes are found up-regulated. In addition, heat-shock protein 90 (HSP90) known as the stressed response gene, and Gr10, the amino acid response gene were up-regulated and down-regulated respectively in the temperature-stressed group. Low expression of Gr10 under temperature-stress can affect nursing behavior and bee development. Ultimately, these findings will help in identifying honeybee-temperature survival mechanisms under varying temperature effects.

## 1.0 Introduction

Recently, there has been an increase in interest in mass colonies and the pollination services that honey bees provide [1,2]. The development of honey bees however is influenced by many factors [1] such as environmental temperature [3]. Abiotic factors especially environmental temperature can affect the internal and external activities of honey bee colonies, including honey bee development, behavioral performance, susceptibility to disease, and production [4–7]. In addition, honey bees have been declining due to colony collapse disorder (CCD), which is also a massive contributing factor to the abrupt disappearance of honey bee colonies [8,9]. CCD has caused severe negative impacts due to the decline of pollination [10]. CCD mechanism remains unknown, although other possible factors such as pesticides, infectious pathogens, genetic factors, and temperature can be considered [11,12].

Climate change appears to be one of the major factors towards temperature changes and a major concern for agriculture and beekeeping [13]. It is also one of the possible contributors to the decline in pollinators, including honey bees [6,14]. Weather-related impacts like temperature intensity have been connected with the activity of social insects such as honey bees [15,16]. Higher air temperatures elevate colonial net gain rates and increase the efficiency of honey storage rates by lowering metabolic rates [17]. Ambient air temperature also affects the honey bee flight activity [18]. However, little detailed information is available under which conditions of weather have an impact on honey bee mortality, and social behaviors [19,20].

Hypopharyngeal glands (HGs) in honey bees are paired glands that develop as bees develop [21]. They are exclusively located in the head region of worker bees and play a significant role in various social behaviors through the production of different secretions [22,23]. Each gland consists of small oval bodies called acini that are connected to terminal or axial secretory ducts. The HGs usually become fully developed in young workers (6-13 days) with a large functional secreting acini [24,25]. Once the bee starts foraging behavior the glands degenerate [26]. The bee development affects gland activity from producing the primary royal jelly (RJ) proteins in young nurse bees to manufacturing carbohydrate-metabolizing enzymes like α-glucosidase in forager bees [27]. RJ is not just a protein-based food fed to larvae but also helps to produce healthy queens [28]. However, these glands are vulnerable to various stresses such as *Varroa* infestation and heat, which may result in the reduction and degeneration of the gland, and subsequently their biological and genetic functions [29].

Heat stress is one of the major factors affecting honey bee mortality, however, they evolved a series of measures to cope with temperature stress such as morphological changes [13]. Summer foragers exposed to extreme heat stress show less activity of HGs compared to nurses who are not exposed to the same stress [30]. The reduced HGs activity resulted in a low protein synthesis rate or decomposition[31], slowing down or repairing the damage by increasing the content of protective substances [32,33]. Heat shock proteins (HSPs) play a vital role in the stress responses of many organisms [34] and act as molecular chaperones to protect the protein from damage during protein synthesis, and folding [34,35]. HSPs of various sizes and functions exist including HSP60, HSP70, and HSP90 [36]. While HGs are fully involved in nursing behavior, other proteins such as GR10 (Gustatory receptor 10) protein that is primarily involved in nursing behavior and other division of labor needs to be examined equally under heat stress. Overall, heat stress triggers the expression of certain genes other than HSPs, that are involved in some biological processes including metabolism, and translation [37]. Surprisingly, little research has been performed on the relationship between climate change and the genetic functions of honey bees concerning HGs performance [38–41]. Especially for it is a nutritional function related to nursing behavior, and biological adaptation, especially under various unfavorable conditions such as temperature and or chemical stress.

Therefore, we compared the transcriptome of *Apis mellifera* HGs at Low temperature, High temperature, and regular temperature using RNA sequencing technology and detected the sequence pattern of some differentially expressed genes (DEGs) in hypopharyngeal tissues respectively. Our data aim at comparing the DEGs from one another among the three conditions to better understand the regulatory mechanisms including the effect of temperature-stressed genes from the DEGs in honey bees’ physiological and biological activities. This study unveils the status of HGs under temperature-stress, and its implication in honey bee physiological and behavioral activities. Our result identified important temperature-stressed genes and canonical pathways that affect HGs’ biological activities and behavior.

### 2.0 Methods

### 2.1 Raring condition of Apis mellifera

The *Apis mellifera* used in this study were bred at the Incheon National University Apiary, a honey bee breeding facility equipped with an enclosed space designed to maintain a stable temperature. The average monthly temperature was regulated at ± 3°C to investigate the physiological response of bees to climate change factors. To monitor the environmental conditions inside the bee colony, temperature/humidity sensors, and a carbon dioxide sensor were installed. Despite fluctuations in the external temperature, the temperature inside the bee colony was consistently controlled at 32-34°C **(Table S1).** Newly born eggs from queen bees adapted to the temperature range (room temperature and monthly average temperature ± 3 °C) were collected for subsequent experiments. These eggs were transferred to an acrylic cage (W50cm × L12cm × H25cm) and incubated at 34°C until eclosion and adulthood. The hypopharyngeal glands (HGs) were dissected, and the survival proportion was quantified **(Figure S1)**. The development stage was properly monitored, honey bees were subjected to the three temperature conditions at the pupa to adult stages **(Figure S2),** and RNA was extracted. The acini size was majored and showed a significant change as well **(Figure S3).** All experiments were repeated three times in June, July, and September 2020.

### 2.2 RNA extraction and cDNA synthesis

RNA was extracted from dissected HG at -75°C to -80°C using dry ice to prevent denaturation by RNeasy® Mini Kit (QIAGEN, Germany) according to the manufacturer’s protocol. cDNA was synthesized from 1 µg of RNA. First, DNase treatment of RNA samples before RT-PCR was performed by RQ1 RNase-Free DNase (Promega, WI, USA). Then reverse transcription was performed with oligo dT with Superscript III enzyme (Invitrogen, CA, USA). The results of RNA extraction of hypopharyngeal glands were presented in **(Table S2).**

### 2.3 RNA sequencing (RNA-seq) and quality control

The total RNA of the nine samples was prepared at 2 µg per sample before experiments. RNA Quality control was examined with 2100 Expert Bioanalyzer (Agilent Technologies, CA, USA) using mirVana ^TM^ miRNA Isolation Kit (Ambion, TX, USA), and RNA 6000 Nano Kit (Agilent Technologies, CA, USA) according to the manufacturer’s protocol. Once all samples passed, library preparation was performed by Truseq Stranded mRNA Library Prep kit (Illumina, CA, USA). Sequencing was performed on the Novaseq 6000 (Illumina, CA, USA) in 100bp PE configuration according to the manufacturer’s instruction. Quality control of raw reads was performed through FastQC [42]. Adapter sequences were trimmed out and low-quality reads were discarded using Trim Galore [43].

### 2.4 Statistical Analysis

The statistical analyses were conducted for developmental stages, acini size, and survival rate. Kaplan-Meier analysis and post-hoc log-rank test were used to derive and compare overall survival curves and temperature variation curves between the groups. Multivariate analysis for survival was performed using the Cox-Mantel model. A p-value of <0.05 was considered to indicate statistical significance. Graphs were created using GraphPad Prism software (version 7.03).

### 2.5 Differentially expressed genes (DEGs) analysis

Gene expression level (FPKM value) was measured with CLC Genomics Workbench 11.0 (QIAGEN, Germany) using RNA-seq analysis tools. Fold change value and p-value were analyzed by calculating the Normalized FPKM value using the Differential Expression for RNA-Seq tool. EnhancedVolcano () R-package was used to generate a volcano plot for significant DEGs, Pheatmap () for heatmap construction, and various plugins for KEGG pathway enrichment analysis.

### 2.6 Functional analysis

GO (Gene Ontology) terms including BP (Biological process), CC (Cellular component), and MF (Molecular function) as well as KEGG (Kyoto Encyclopedia of Genes and Genomes) pathway ID were identified using DAVID 2021 tool [44–46]. The genes were ranked by the log_2_FC value of RNA-seq data, and Venny 2.1 was used to identify the commonly shared genes between the three groups. ggplots () R-package was used to present the GO-terms.

### 2.7 Construction of Protein-Protein Interaction Network

The protein-protein interaction (PPI) network was constructed using the STRING v11.5 database plugin from Cytoscape v3.9.1 for network analysis. Cytoscape plugins including CytoHubba v0.1 for the identification of highly connected genes (Hub-genes) based on their degree of connectivity. The MCODE v2.0.2 was utilized to identify interconnected regions or clusters from the PPI network, The cluster finding parameters were adopted, such as a degree cutoff of 2, a node score cutoff of 0.2, a kappa score (K-core) of 5, and a max depth of 100, which limits the cluster size for the co-expressing network. ClueGO v2.5.9 was used for detailed functional annotation among the DEGs.

### 3.0 Results

### 3.1 Survival Rate and Development

Under variable temperatures, there is a negative survival rate of the HGs tissue when exposed to higher temperatures. On the other hand, a positive survival rate was observed at lower temperatures and average with a significant value of p<0.043. The acini size checked in adults shows less development when subjected to a high temperature as compared to the other two conditions. This might indicate the effect of high heat stress on acini development. This result remains the same bot in pupa and adult honey bees’ HGs. Overall, high temperature affects the honey bee development as well as survival rate.

### 3.2 Identification of DEGs and top-regulated genes

A differential expression gene (DEG) analysis was conducted comparing Low temperature with High temperature, Regular temperature with High temperature, and Regular temperature with Low temperature, respectively. Volcano plots for the DEGs showed different genetic profiles between each group (**Figure 1A-C**). In addition, the Venn diagram represents the combination of DEGs (2Fold change), with 5196 identified as commonly expressed genes (**Figure 2A**). DEGs from the three groups indicated more down-regulated genes compared with up-regulated genes (**Figure 2C**). There was variation among the top 10 DEGs from each group which include *LOC100577410, LOC100576982, and OBP13* as top up-regulated genes and *LOC561044, LOC724239, and LOC409708* among the top down-regulated genes for the Low Vs High group. Also, *LOC113219378, LOC113218757, and LOC412162* as top up-regulated genes, and *Apd-3, LOC102653731, and LOC408614* were among the top down-regulated genes for the Regular Vs High group. For the Regular Vs Low group, *LOC113219378, LOC113219355, and LOC552224* were among the top up-regulated genes, while *LOC726850, LOC100577092,* and *LOC113219358* were among the top down-regulated genes as shown in (**Figure 2C**).

**Figure 1.**
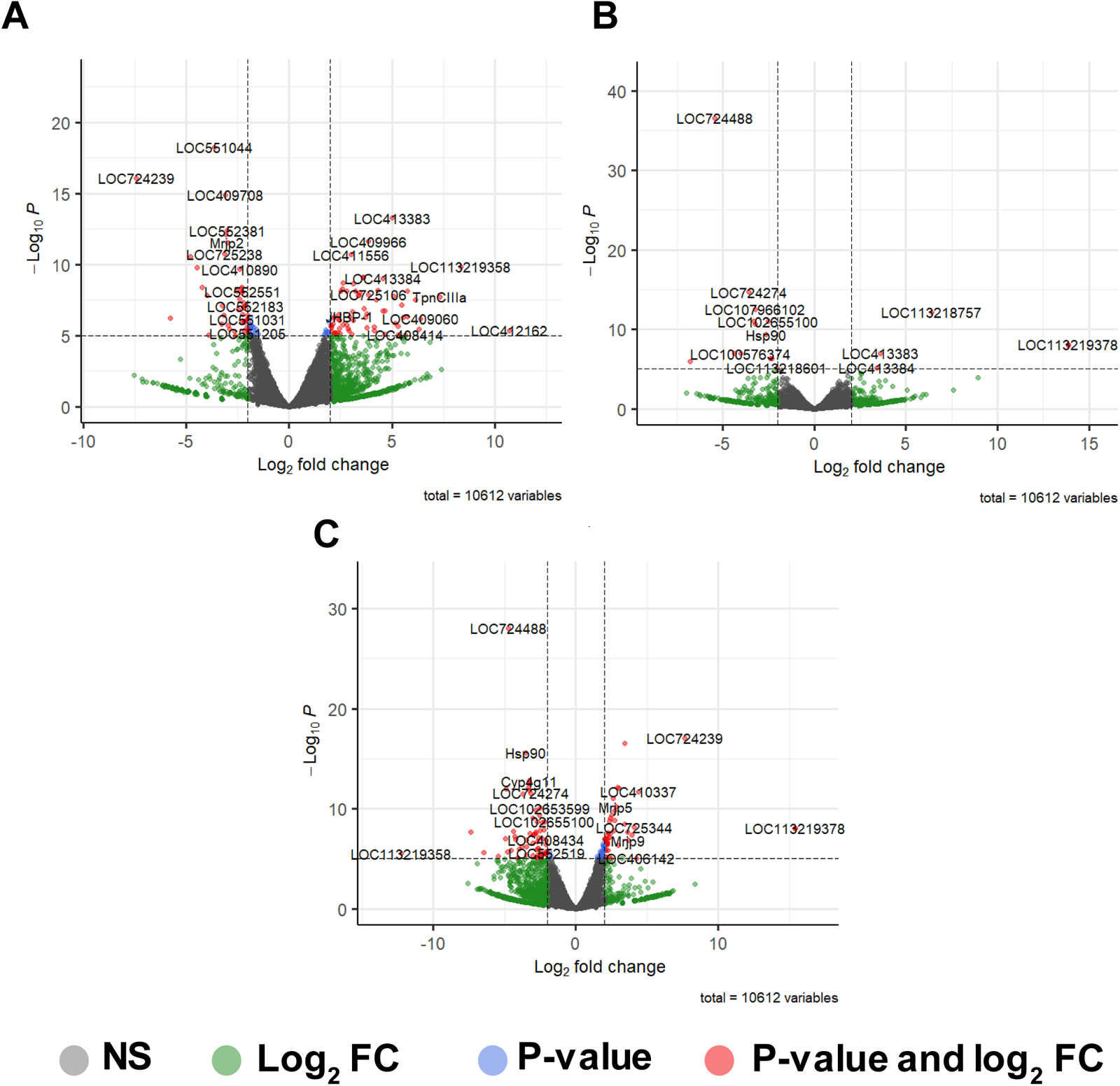
**(A)** Enhanced volcano plot of differentially expressed genes identified between the Low-temperature group and High-temperature group. **(B)** Enhanced volcano plot of differentia l ly expressed genes identified between the Regular temperature group and High-temperature group. **(C)** Enhanced volcano plot of differentially expressed genes identified between the Regular-temperature group and Low-temperature group. Green color dots indicate differentially expressed genes (DEGs) based on Log_2_ fold change. Blue color dots indicate significant genes based on the p-value. Pink color dots indicate genes with p-value and log_2_ fold change (FC). Genes with positive log_2_ FC are up-regulated, while genes with negative log_2_ FC are down-regulated.

**Figure 2:**
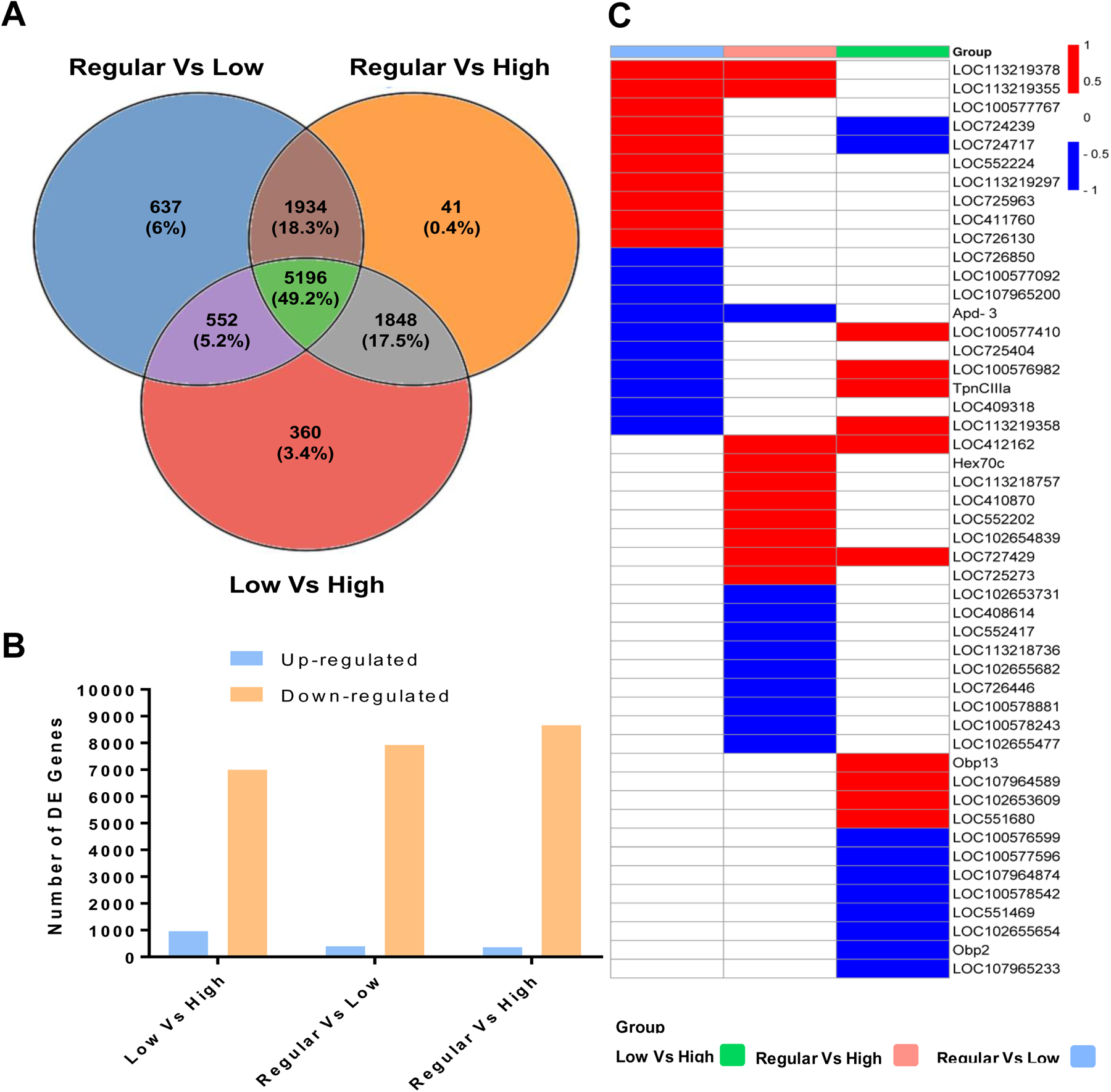
**(A)** Venn diagram representing the combination of the differentially expressed genes (DEGs) from the three groups. **(B)** Bar graph showing the number of Upregulated and down-regulated DEGs from the three groups. **(C)** Heatmap showing the expression pattern of the top then (10) DEGs from each group. Red color; Up-regulated, Blue color; Down-regulated, and White color, not present.

### 3.3 Functional Enrichment Analysis for Common DEGs

Database for Annotation, Visualization, and Integrated Discovery (DAVID) database was used to annotate the 5196 common DEGs based on their BP, CC, and MF. The top five (5) GO terms were represented including RNA processing (GO: 0006396), RNA metabolic process (GO: 0016070), and protein modification process (GO: 0036211) for BP. CC GO terms include nucleoplasm part (GO: 0044451), nucleoplasm (GO: 0005654), and Golgi membrane (GO: 0000139) among others. The top five (5) enriched MF include RNA binding (GO: 0003723), ribonucleoside binding (GO: 0032549), and purine nucleoside binding (GO: 0001883) among others (**Figure 3A**). KEGG was analyzed, and the top ten (10) enriched pathways were sorted out including glycerophospholipid metabolism (ame00564), nucleocytoplasmic transport (ame03013), ribosome biogenesis in eukaryotes (ame03008), and MAPK signaling pathway – fly (ame04013) among others (**Figure 3B**).

**Figure 3:**
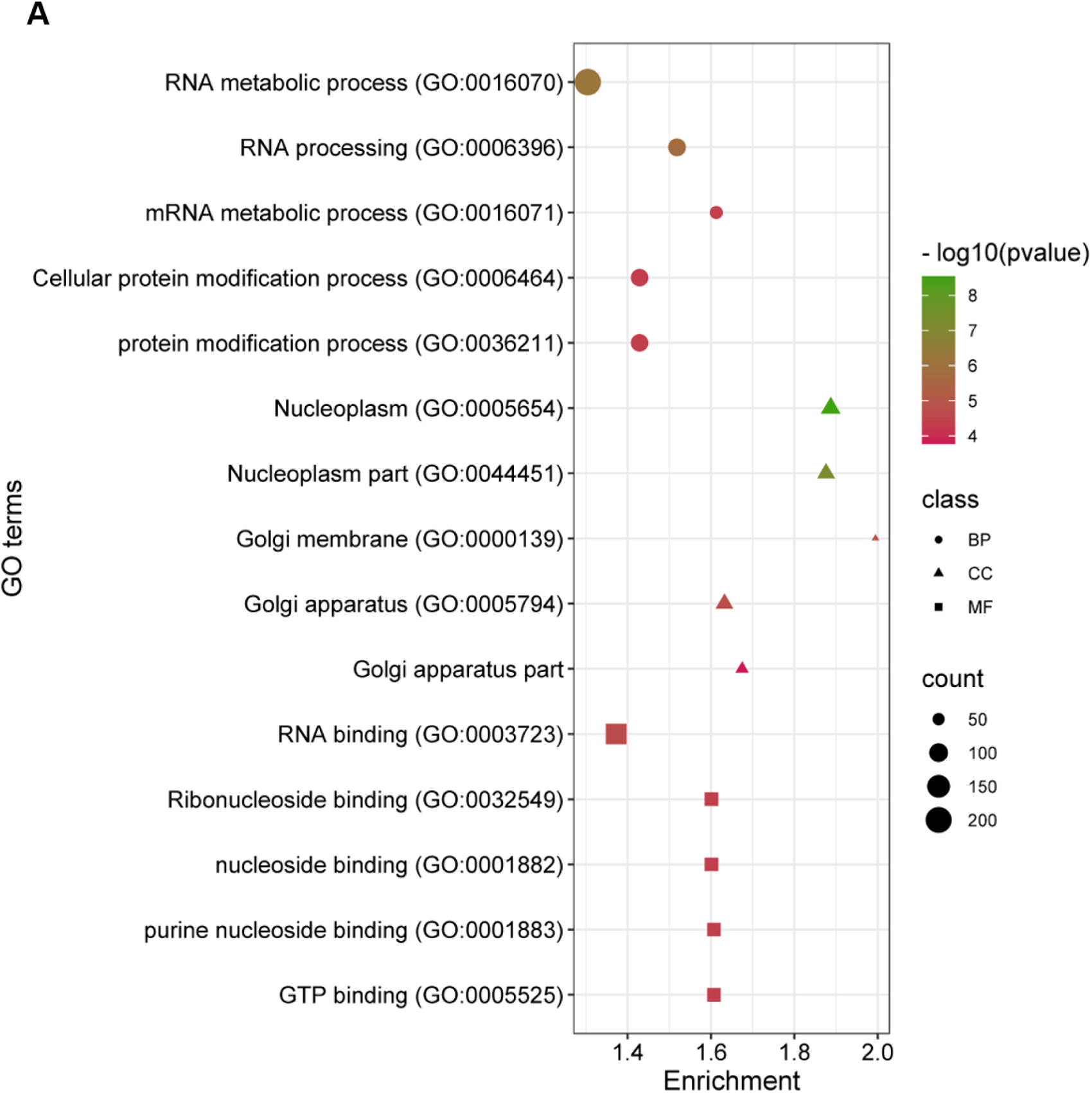
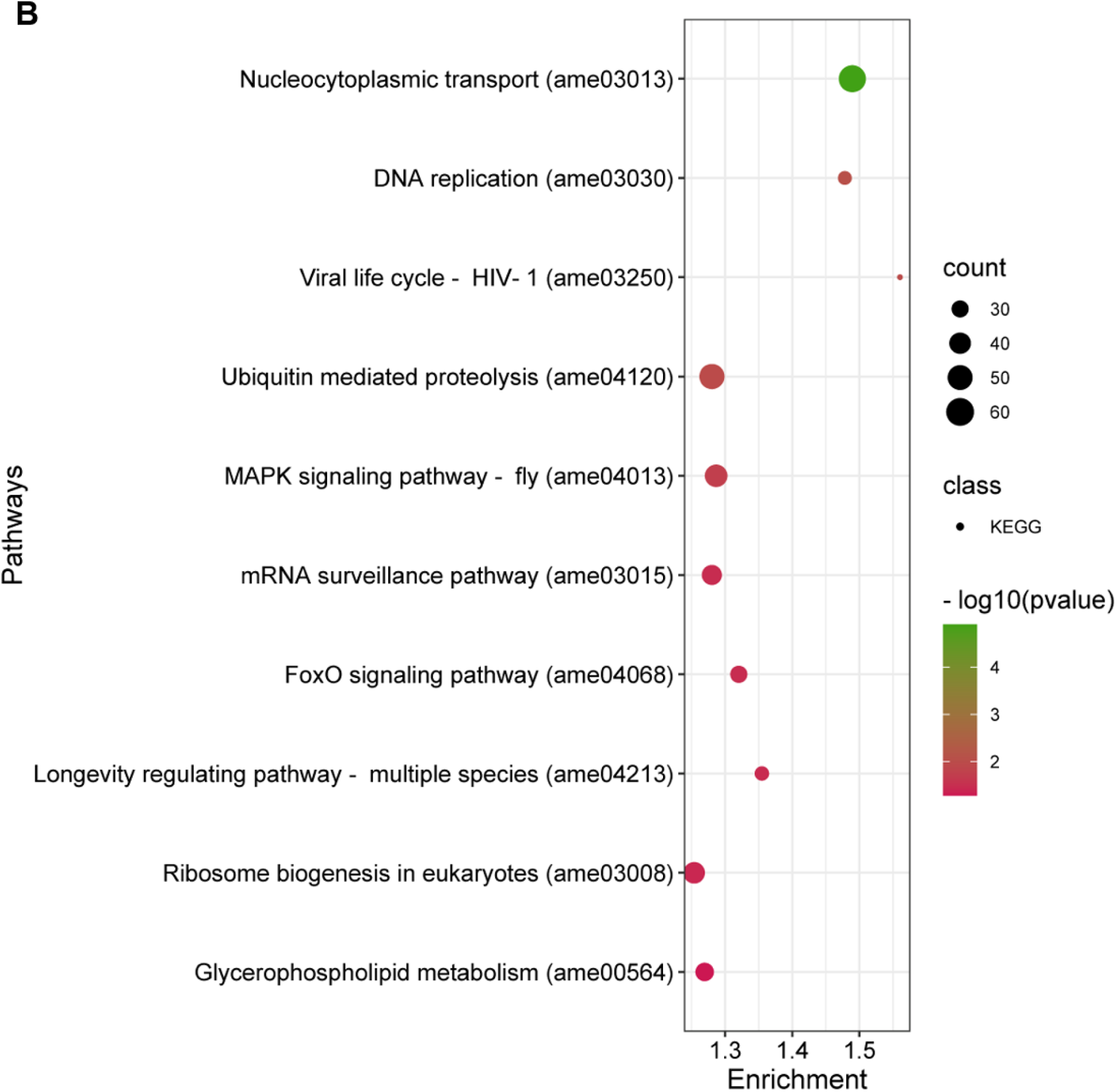
**(A)** Bubble chart indicating the top five (5) gene ontology (GO) enriched terms for the commonly differentiated genes (cDEGs). This includes BP; Biological process. CC; Cellular component. MF; Molecular function. Significant GO terms are based on the number of genes involved. **(B)** Bar chart for KEGG pathway enrichment analysis of the cDEGs. Only the top ten (10) were included.

### 3.4 Protein-Protein Interaction Network and Enrichment Analysis for Common Genes

Further functional enrichment analysis was performed using the cDEGs for their PPI (protein-protein interaction). To comprehensively dissect the GO terms and the pathway associated with these genes, the ClueGO plugin from Cytoscape was used. Highly enriched GO terms included adenyl ribonucleotide binding, nuclear part, protein modification by small protein conjugation or removal, ATP-dependent helicase activity, regulation of cyclin-dependent protein serine/threonine kinase activity, regulation of RAS protein signal transduction among others, and RNA processing (**Figure 4A**). The plugin provides an option specifically for highly enriched KEGG pathways. The identified highly enriched pathways include d ribosome, oxidative phosphorylation, and citrate cycle (TCA cycle) (**Figure 4B**). The CytoHubba plugin from Cytoscape software was used to identify the top ten (10) hub genes based on their degree of connectivity. These include *LOC413462* as the highest node followed by *GB41233-PA, GB53954-PA, LOC413935, LOC413404, LOC4109590, LOC410907,* and *LOC724129* (**Figure 4C**). Three of these hub genes were found to be highly enriched in pathways related to ribosome biogenesis in eukaryotes. MCODE plugin v2.0.2 from Cytoscape was utilized to identify the densely interlinked regions within the PPI. As a result, we obtained the top three (3) significant clusters/modules from our common DEGs PPI network with scores of 62.54, 15.15, and 14.07 for Cluster 1, Cluster 2, and Cluster 3 respectively (**Figure S5)**. For each cluster, a bottleneck gene was identified. These included *LOC412931, LOC726336,* and *LOC409340* for Cluster 1, Cluster 2, and Cluster 3 respectively. To provide more detailed information about this sub-network, their enriched KEGG pathway was identified which includes ribosome biogenesis in eukaryotes (Cluster 1), valine, leucine and isoleucine degradation, fatty acid degradation, and propionate metabolism among others (Cluster 2), while spliceosome was identified as the most enriched pathway for (Cluster 3).

**Figure 4:**
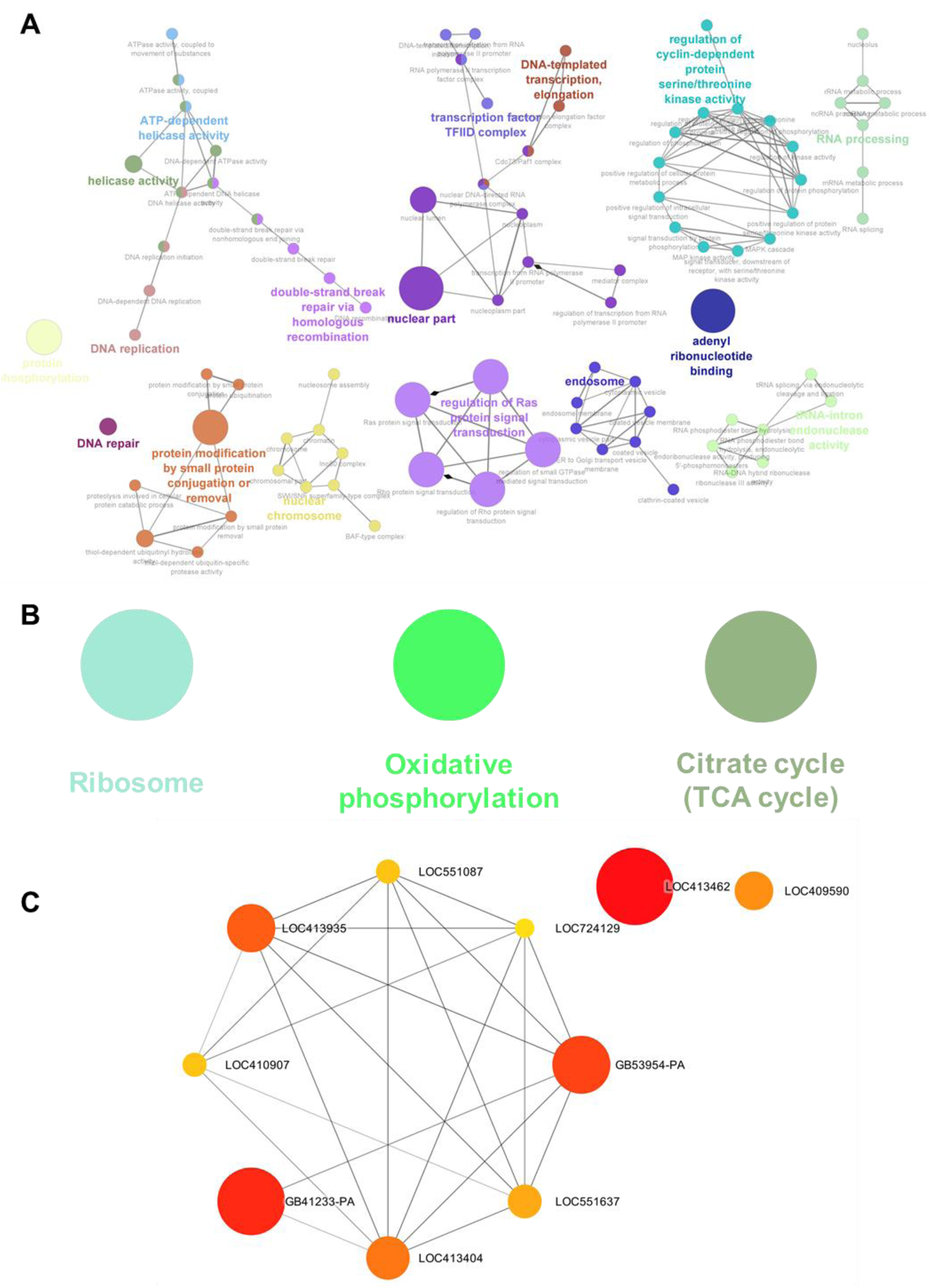
**(A)** Cytoscape-based ClueGO visualization of highly enriched GO-terms including BP; Biological process. CC; Cellular component. MF; Molecular function. Only the most significant terms are included. The bigger the node the higher the enrichment. **(B)** Cytoscape-based enrichment analysis and visualization of the cDEGs. The enriched pathways were derived from the Kyoto Encyclopedia of Genes and Genome (KEGG) database. Three highly enriched pathways were reported. Terms are linked based on к-score (≥0.4), edge thickness indicates the association strength while node size corresponds to the statistical significance for each term. **(C)** Top ten (10) hub genes identified from the cDEGs protein-protein interaction (PPI) network. The bigger the node size the higher the degree of connectivity.

### 3.5 Functional Analysis for Temperature-Stressed Genes

Temperature-stressed genes were unique DEGs identified from the Low Vs High-temperature group. The Venn diagram from **Figure 2A** identified 360 unique genes, among which 118 are up-regulated, while 242 are down-regulated **(Table S3).** DAVID online tool was used to identify their enriched GO terms including BP, CC, and MF as well as KEGG. The top ten (10) enriched BP include translation, peptide biosynthetic process, amide biosynthetic process, and peptide metabolic process among others. Enriched CC includes ribosome, intracellular ribonucleoprotein complex, and ribonucleoprotein among others. Enriched MF included structural constituent of ribosome, structural molecule activity, and RNA binding among others (**Figure 5A**). KEGG pathways such as the ribosome, fatty acid metabolism, citrate cycle (TCA cycle), and oxidative phosphorylation were identified as the enriched pathways (**Figure 5B**).

**Figure 5:**
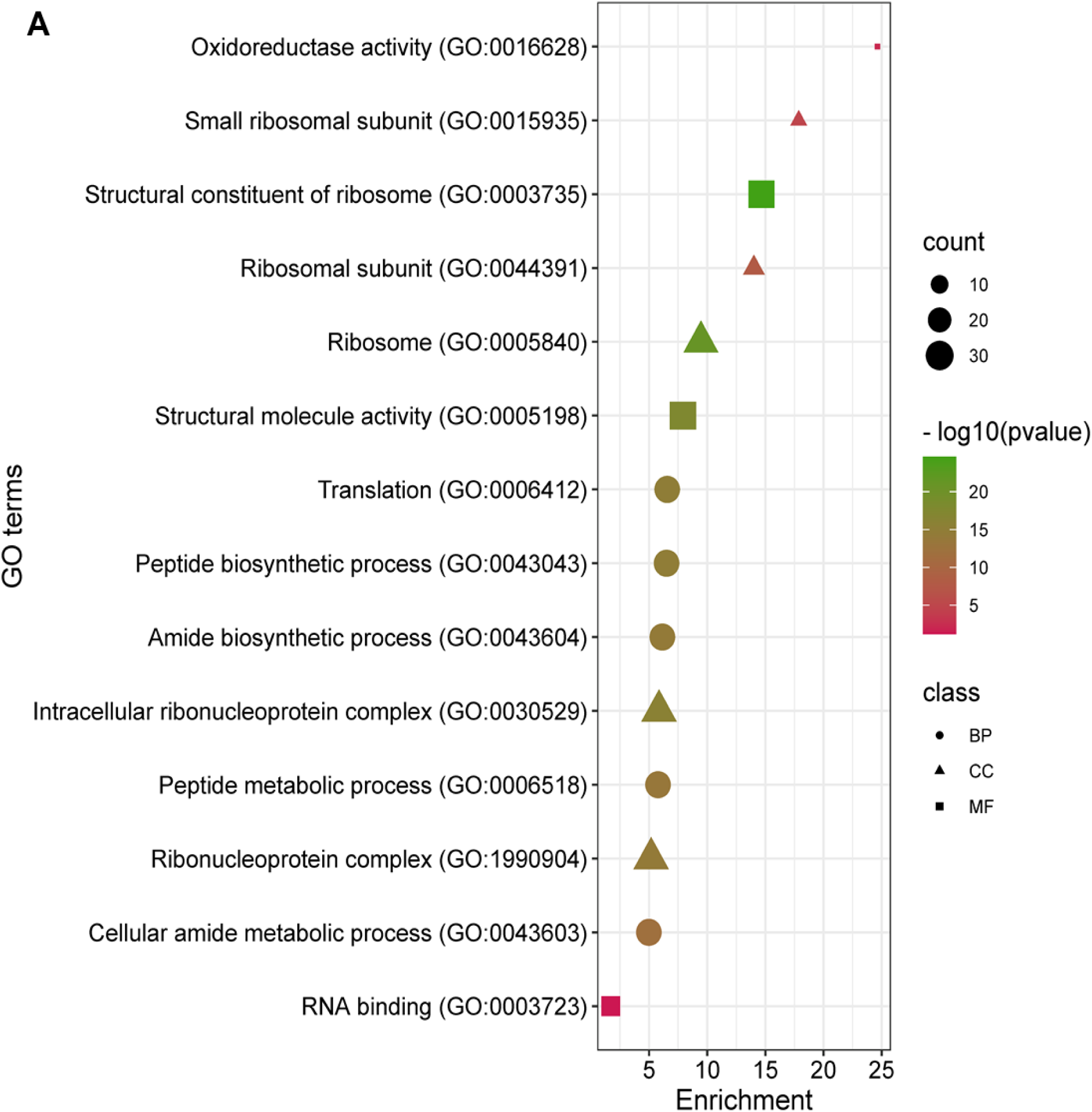
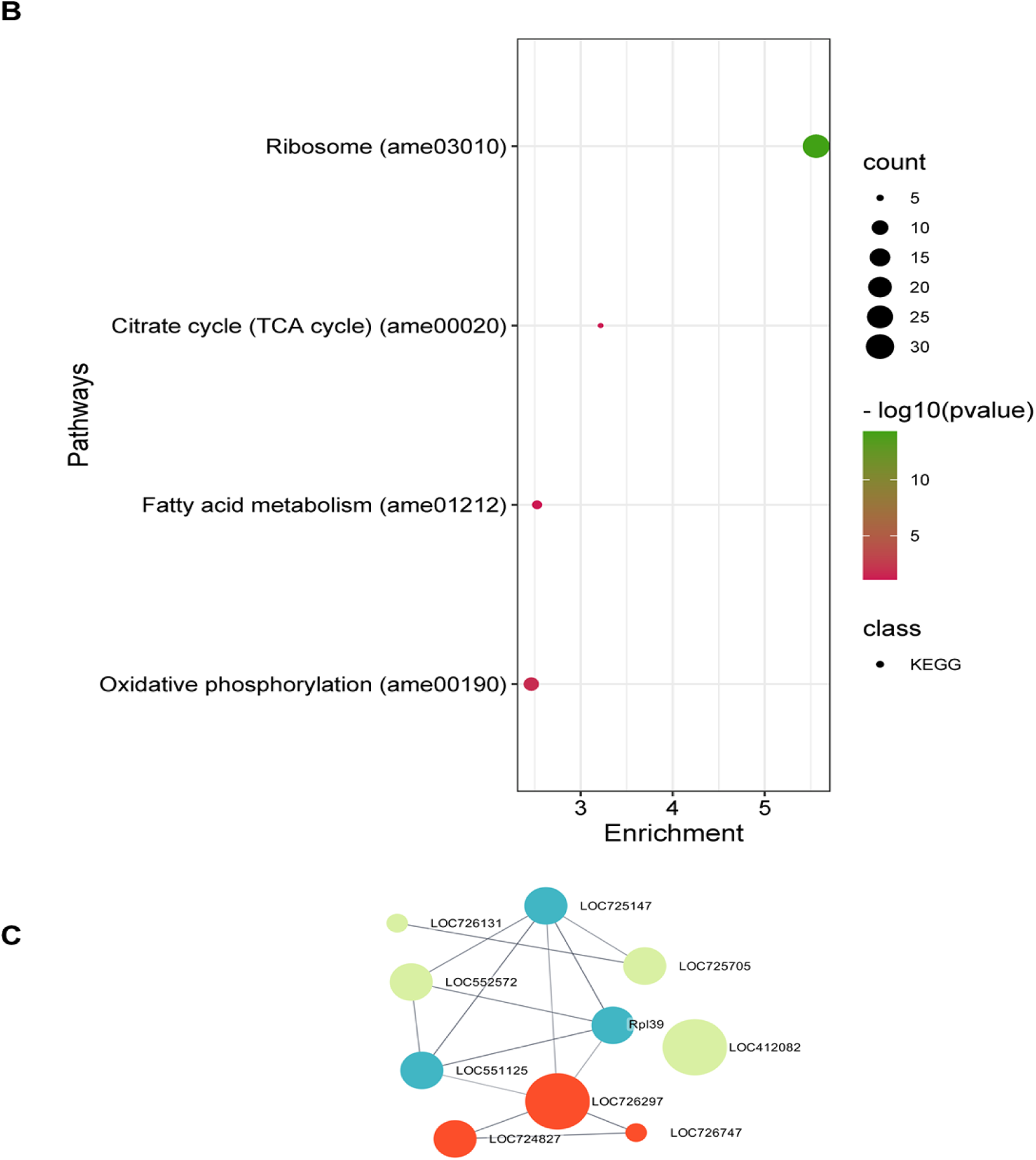
**(A)** Bubble chart indicating the top ten (10) enriched gene ontology (GO) for the temperature-stressed genes. GO-terms include Biological process, Cellular component, and Molecular function. **(B)** Bar chart for KEGG pathway enrichment analysis of the identified temperature-stressed genes. **(C)** Cytoscape-based analysis using CytoHubba identified the top ten (10) hub genes identified from the temperature-stressed genes protein-protein interaction (PPI) network. The bigger the node size the higher the degree of connectivity.

**Table 1:** List of top 10 hub genes and their expression pattern under temperature-stressed conditions.

| Gene symbol | Log2 FC | p-value | FDR p-value | Expression |
| --- | --- | --- | --- | --- |
| LOC726297 | -0.37195255 | 0.393081571 | 0.638282002 | Down-regulated |
| LOC412082 | -0.310074289 | 0.570563228 | 0.773296415 | Down-regulated |
| **LOC551125** | **0.130230254** | **0.722380878** | **0.864090075** | **Up-regulated** |
| **LOC724827** | **0.248892926** | **0.53523573** | **0.749619137** | **Up-regulated** |
| LOC725147 | -0.231282146 | 0.614853285 | 0.800797569 | Down-regulated |
| LOC725705 | -0.925793392 | 0.13013334 | 0.341080024 | Down-regulated |
| LOC552572 | -1.01410744 | 0.010898236 | 0.071348223 | Down-regulated |
| Rpl39 | -0.14915706 | 0.765181762 | 0.88658128 | Down-regulated |
| LOC726131 | -0.012880032 | 0.989167512 | 0.994946035 | Down-regulated |
| LOC726747 | -0.452448322 | 0.302694404 | 0.551362964 | Down-regulated |

### 3.6 Enrichment Analysis and Identification of Temperature-Stressed Hub Genes

The PPI network for 360 temperature-stressed genes was constructed and the top 10 hub genes were identified using the CytoHubba plugin from Cytoscape. Among the 360 DEGs, 10 hub genes were identified including *LOC412082, LOC726297, LOC724827, LOC725147, LOC551125, LOC552572, LOC725705, RPL39, LOC726131,* and *LOC726747* (**Figure 5C**). From the DEGs analysis data, all the top 10 genes were found up-regulated in the Low Vs High-temperature group. After mapping them with the KEGG database, the hub genes are enriched in the ribosome and oxidative phosphorylation pathways. Specifically, only three genes are involved in each pathway including *RPL39, LOC725147,* and *LOC551125* for the ribosome pathway, while *LOC726297, LOC724827,* and *LOC726747* for the oxidative phosphorylation pathway.

## 4.0 Discussion

Honey bee colonies recently faced many challenges, which include both biotic and abiotic factors. The impact of abiotic factors specifically temperature on honey bees has been previously reported [18,47,48 ]. The normal temperature range within a honey bee colony ranges from 22 to 36 °C [49,50]. On the individual level, the temperature rises above 36 °C lead to honey bees exposing their brood to overheating, and induced HSPs to resist any passive effect [51]. An increase in the level of HSPs was observed among the worker bees exposed to a variable temperature between 4 to 50 °C [52], highlighting the impact of temperature on the bee genetic changes. Generally, organisms that are able to rapidly alter gene expression have an evolutionary advantage, which is crucial for their survival [53]. Therefore temperature adaptation often involves various transcriptional regulatory processes [54].

The honey bee workers possess HGs which play an important role in their nutrition [55]. The HGs in honey bee workers undergo a transition through undeveloped, developed, and regressive phases, coinciding with their shift in tasks from nursing to foraging within a 20-day timeframe in a typical colony. In young workers, typically within the first 13 days post-eclosion, the HGs, which are fully developed and exhibit robust protein synthesis during the nursing phase, begin secreting RJ for brood rearing. As workers reach the 18th day and transition into foragers, the HGs start to regress and produce enzyme products to facilitate the digestion of nectar into honey [28,56]. Previous research focused mainly on morphological, physiological, biochemical, age-dependent, and proteomics network analysis to follow up protein changes during the HG development under natural conditions [55]. However, subjecting the HG to various stresses such as temperature might lead to significant protein changes that can be related to it is physiological, morphological, biological changes, and importantly nutrition. In the past, genetic relationships and adaptation to the climatic conditions of *Apis cerana* based on genetic polymorphism have been established [57]. Our study mainly explored the effect of temperature on honey bees (*Apis mellifera*) HGs under three conditions including low, high, and regular temperature.

In this study, we identified DEGs between the honey bees’ HGs treated under three temperature conditions. Our data and protocol are in line with the existing research on other insects. In the past, RNA-seq in New Zealand stick insects showed that genes expressed under cold temperatures (-5 °C) included cuticular genes and other genes related to physiological variation and carbohydrate metabolism [58]. Proteomics and transcriptomic analysis in Chinese white wax insects (*Ericerus pela*) from various climatic regions found 2386 DEGs, involved in various biological processes including catalytic activity, and response to stimuli [59]. From our DEG analysis, a total of 5196 common DEGs (cDEGs) identified occupied 49.2% of the total DEGs combined. Down-regulated genes are significantly higher than up-regulated genes in all three groups. Next, based on GO analysis, the cDEGs were enriched in BP specifically RNA processing, and mRNA metabolic process among others. It is not surprising that the detected genes are involved in the metabolic process as the main energy source of the honey bee is glucose, fructose, sucrose, and maltose [60,61].

The pathway analysis revealed that the cDEGs are highly enriched in Glycerophospholipid metabolism, nucleoplasm transport, MAPK signaling, and FoxO signaling pathways as the most significant pathways. The primary driving metabolites in honey bees are reported to be highly involved in Glycerophospholipid metabolism and are equally up-regulated in the brain [62]. This highlights Glycerophospholipid metabolism’s critical role in brain physiology. Also, thiacloprid used as an insecticide was found to cause behavioral dysregulation in honey bees by significantly affecting the Glycerophospholipid metabolism pathway [63], indicating the impact of xenobiotic stress on bee metabolism. Asymmetric glycophospholipids allow cells to adapt to temperature variations and hypoxic environments in yeast [64].

Highly connected cDEGs were rated based on their degree of connectivity. The top 10 highly connected nodes called hub genes include *LOC413462* (Arachidonic acid metabolism), and *GB41233-PA* among others are the key nodes for the HGs response to temperature variations and are highly enriched in ribosome biogenesis in the eukaryote pathway, oxidative phosphorylation, citrate cycle (TCA cycle), and fatty acid metabolism among others. Thermal stress results in alterations in genes that are involved in ribosome biogenesis, and various metabolic pathways [65]. Oxidative phosphorylation on the other hand is an essential phenomenon for energy generation. A decrease in oxidative phosphorylation activity causes increased aggression in honey bees and fruit flies [66]. Similarly, pharmacologically treating honey bees to inhibit oxidative phosphorylation pathway increasedhoney bee aggression [66]. Lalouette et al [67] reported that cold stress causes oxidative damage in insects, and a warm recovery period activates the antioxidant system while repairing the cold-induced damage. Further, high temperatures in insects perturbed a mitochondrial function which is linked to oxidative phosphorylation [68,69]. This can be extended to the citrate cycle (TCA cycle) oxidative metabolism which also happens within the mitochondria. Both elevated temperatures and pesticide exposure exert a significant influence on oxidative phosphorylation in foragers by inducing alterations in gene expression. [70].

Further, we investigated the detailed enriched GO-terms from the cDEGs including regulation of RAS protein signal transduction (**Figure 4a**). RAS overexpression is reported to activate autophagy [71,72], and promote longevity [73]. RAS/MAPK signaling pathway is not just involved in cellular activities leading to the honey bees’ longevity [74] but also regulates cell survival, and inhibiting cell apoptosis [75]. When ribosome S6 kinase (RSK) is phosphorylated by RAS, it leads to the activation of ribosomal proteins, which in turn facilitates protein synthesis [76]. Proteins such as hemolymph are correlated with dramatic changes in seasons and temperature, altering honey bee physiological activities [77]. Other proteins such as HSPs are essential not only for temperature changes but also facilitates protein synthesis.

The 360 identified temperature-stressed genes derived from Low vs. high-temperature groups, otherwise referred to as temperature stress groups are highly enriched in translation, and peptide biosynthetic process as the most enriched BP GO-terms among others. The enriched pathways derived from the hub genes among the 360 unique genes are similar to the enriched pathway for the cDEGs discussed. The 360 genes are uniquely expressed only in this group. It is well known that the heat-shock response is one of the antagonistic factors in honey bees [78], and is expressed under temperature-stress conditions to neutralize the stress effect [51]. Interestingly, from our data, the heat shock protein 90 (HSP90) expression was down-regulated in two groups (Regular vs. low; Regular vs. high) but up-regulated in the temperature-stressed group (Low vs. high group) as shown in **(Figure S4)**.

The excess temperature of honey bees induced the 70-kDa HSP and their encoding genes [79]. HSPs expressed in nurse and forager bees at various temperatures, although the foragers are reported to be more heat tolerant with the least HSP expression [80]. HSPs primarily serve as molecular chaperones, aiding in the folding of proteins, averting protein aggregation, and guiding misfolded proteins toward specific degradation pathways [81]. In the honey bee, the enhanced expression of HSPs was reported under heat stress conditions [51], which could be a defense mechanism to protect the honey bee against the stress condition. HSPs have been widely identified in worker larvae [82], embryos [83], in head and brain of workers [84], the hemolymph [85], and even the venom gland [86] respectively, and now also in HGs suggesting that they probably act as molecular chaperones by correcting the folding of unfolded proteins usually taking place in the ribosome, thus helping in the cell maintenance and RJ secretory activity of the HGs under temperature stressed condition for honey bee survival and nursing activities.

Further, Neven et al [87] reported that thermal stress is associated with the expression of HSP70 and HSP90. Added to that, it’s evident that high temperature has a significant effect on candidate gene expression (HSP70 and HSP90) not only in HGs but also in the brains of honey bees [88]. It is worth knowing that, HSP90 may promote cell survival and or cell growth [51,89]. A reduction in HSP90 expression under temperature stress might have explained why we identified a negative survival rate as well as low acini size under high temperature. According to the current literature, HSPs in honey bees are induced at high temperature [51], or when the honey bee is stressed under pathogenic infections [90]. Chemical stress in worker bee HGs also induces HSP70 localization in cells and HSP90 activity[91]. Overall, honey bee HSPs are responsive to various proteostatic stresses and potentially promising biomarkers of honey bee stress [92]. While HGs are fully involved in nursing behavior, other proteins such as GR10 proteins that are primarily involved in nursing behavior have been carefully checked. GR10 expression reduced significantly under temperature-stressed conditions. *AmGr10* a GR10 coding gene in *Apis mellifera* is crucial to nursing or brood-caring behavior in the hive and helps explain how the GR protein family in social insects mediates the synthesis of RJ and contributes to behavior in the hive [93]. In addition to its role in nursing behavior, *AmGr10* also has a nutrient receptor function in the HGs, brain, and ovary [93], similar to its counterpart GR43a which senses hemolymph fructose in *Drosophila* [94]. Together, the down-regulation of GR10 under temperature-stress conditions might affect HG’s nursing behavior and honey bee nutrition in general.

Taking physiology and behavior into consideration, temperature affects honey bee colonies as well as foraging activities [95]. The metabolism of honey bees is influenced not just by temperature but also by their exposure to pesticides, which has consequences for the energy metabolism of honey bee foragers. This, in turn, has a detrimental impact on both the olfactory deficits [96] and memory [97] as well as the immune system function and gene expression of honey bees [70]. This phenomenon should equally be applied using the HGs for a clear conclusion.

## 5.0 Conclusion

In conclusion, our results showed that temperature variation affects the biological and genetic functions of HGs in *Apis mellifera* by altering the gene expression patterns. Based on our DEGs analysis, we have identified 360 temperature-stressed genes that are unique. Similar to other findings, our results showed that the expression of HSPs (specifically HSP90) significantly changes in response to various temperature stress, and up-regulated only in the temperature-stressed group but not in the other two groups. Overall, the RAS/MAPK pathway, ribosome, and fatty acid metabolism-related pathway can be highly affected under temperature-stressed which can be related to acini size development, energy development, and survival. In addition to that, GR10 which is responsible for HGs nursing behavior was down-regulated under temperature-stress conditions signifies the negative effect of stress on HG nursing activity, in addition to the significant reduction of the acini size under higher temperature. The impact of temperature on honey bee physiology and behavior often closely resembles the effects of pesticides. Consequently, exploring the honey bee genome to identify crucial marker genes associated with temperature stress could be a promising approach to enhancing their survival, and nursing behavior. Also, silencing *AmGr10* from HG tissue can give us more insight into its genetic role in HG and the honey bee entirely.

## List of abbreviations

DEGs: Differentially expressed genes
cDEGs: Commonly differentially expressed genes
HSP: Heat shock protein
GO: Gene ontology
BP: Biological process
CC: Cellular component
MF: Molecular function
KEGG: Kyoto Encyclopedia of Genes and Genomes
DAVID: Database for Annotation, Visualization, and Integrated Discovery
PPI: Protein-protein interactions
STRING: Search tool for the Retrieval of Interacting Genes
MCODE: The molecular complex detection

## Declarations

### Ethics approval and consent to participate

Not applicable

### Consent for publication

Not applicable

### Authors’ contributions

H.W.K. designed research; A.Y.M., J.H.L., and S.L. performed research; A.Y.M, analyzed data; A.Y.M., Writing-original draft of the paper. H.W.K. J.H.L Writing-review and editing. All authors reviewed the manuscript. The author(s) read and approved the final manuscript.

## Acknowledgments

We thank M. L. Lee and S. Lee for bee colony assistance; J. H. Park in 3BIGs for RNAseq.

---

**Figure S1.**
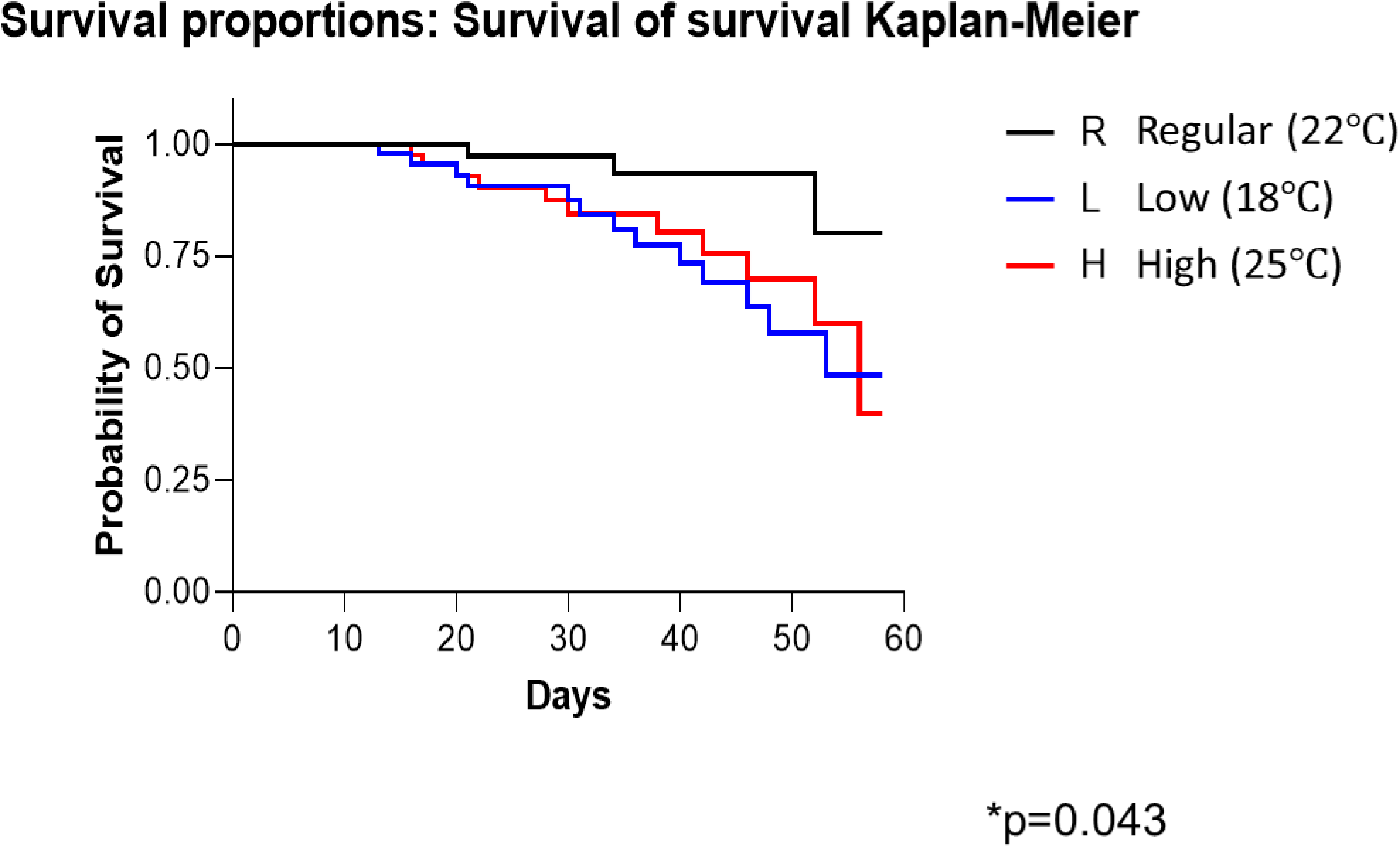
Each honey bee group was compared with a Kaplan-Meier-Survival analysis for survival. A post-hoc Log-Rank test (Cox-Mantel) revealed significant differences between those groups (Log-Rank p=0.043). The blue line indicates positivity, while red indicates negativity. Asterisk indicates statistically significantly lower mortality when compared to the control group.

**Figure S2.**
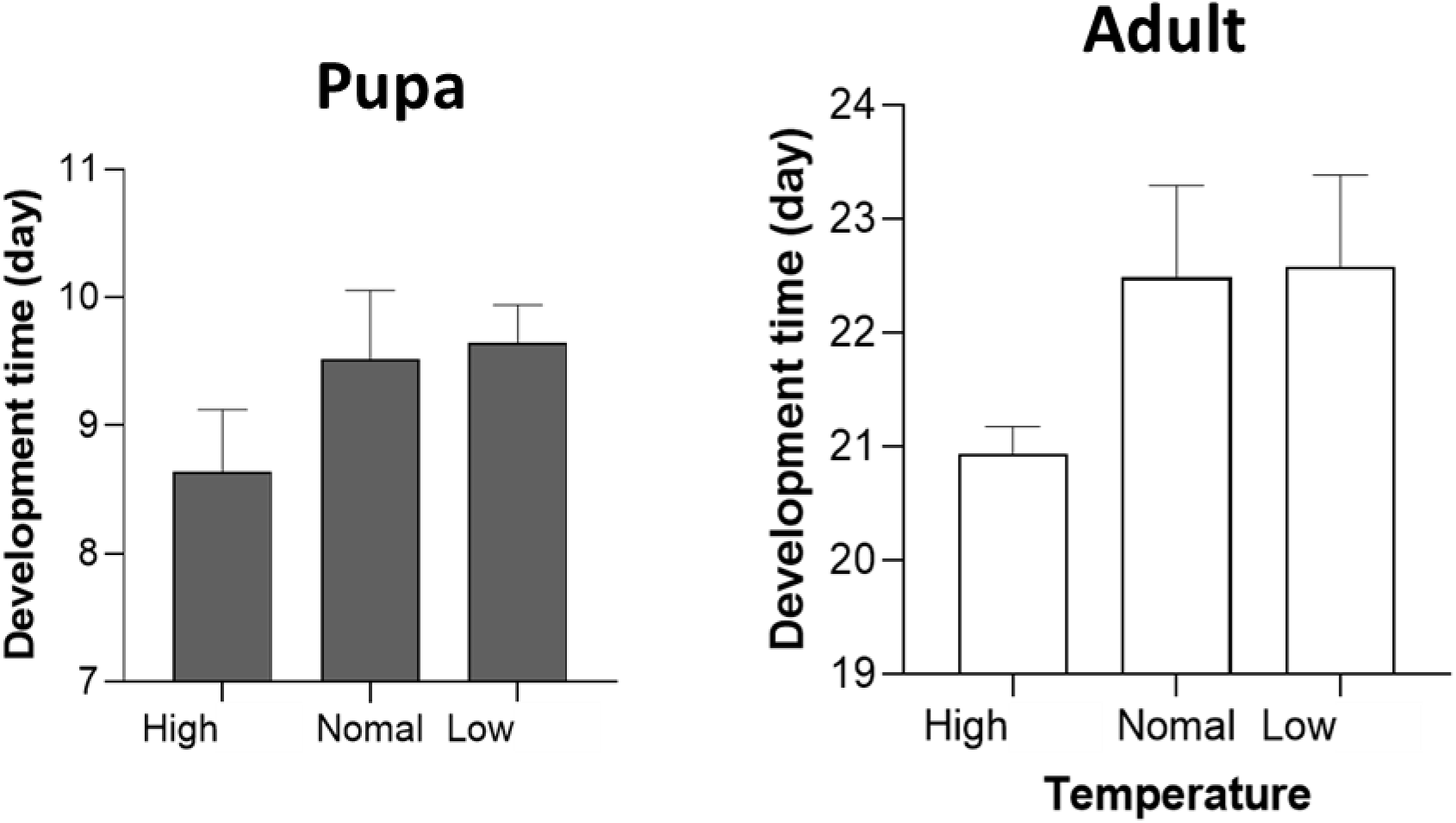
Honey bee developmental stage was monitored from pupa to adult stage under various temperatures. Development timing was rated up to 24 days and 11 days for Adults and pupa respectively.

**Figure S3.**
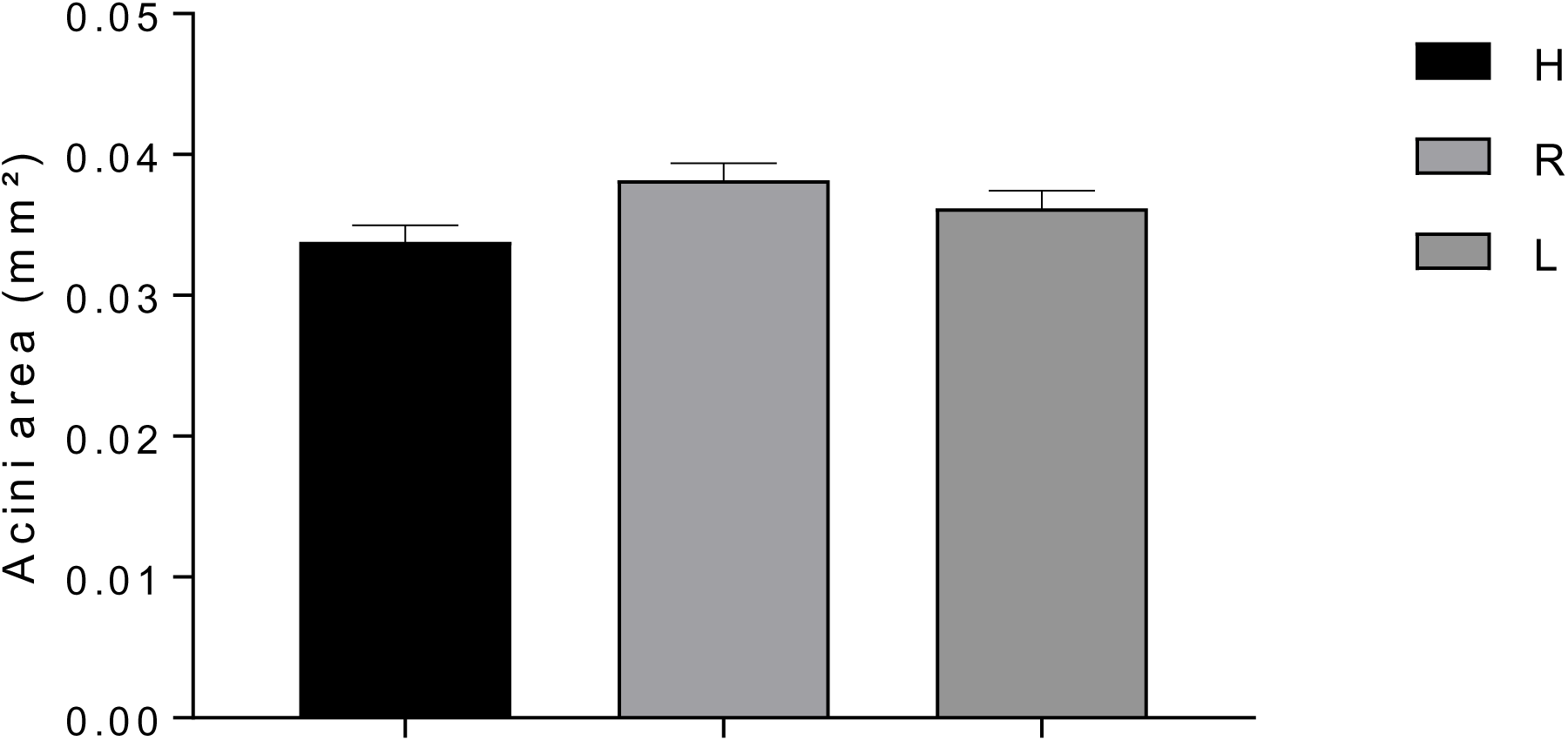
Honey bee developmental stage and acini size show significant change. Where H=High temperature; R=Regular temperature; L=Low temperature.

**Figure S4.**
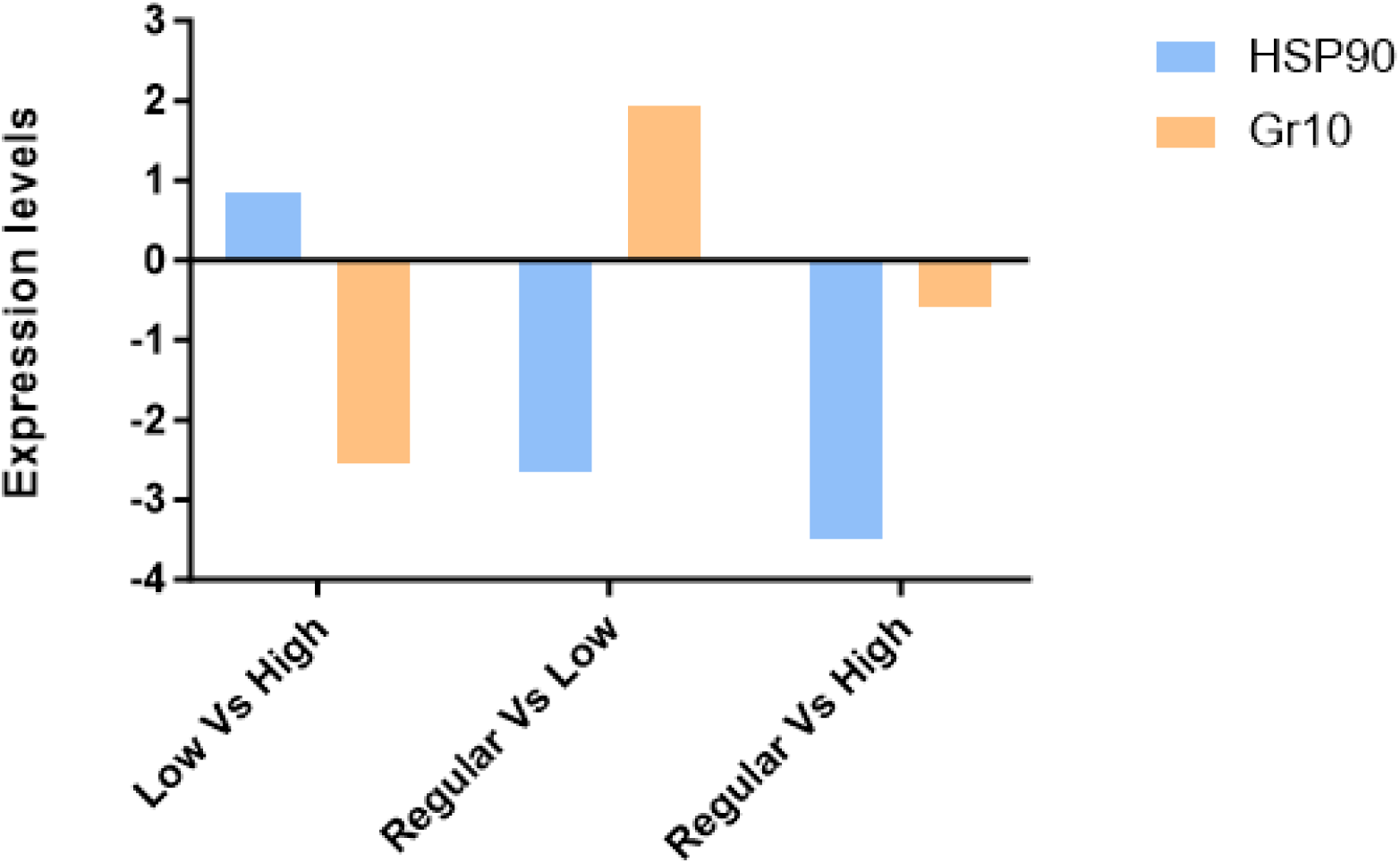
HSP90 and Gr10 expression patterns within the three groups. Low Vs Hight, Regular Vs Low, and Regular Vs High (Temper conditions).

**Figure S5.**
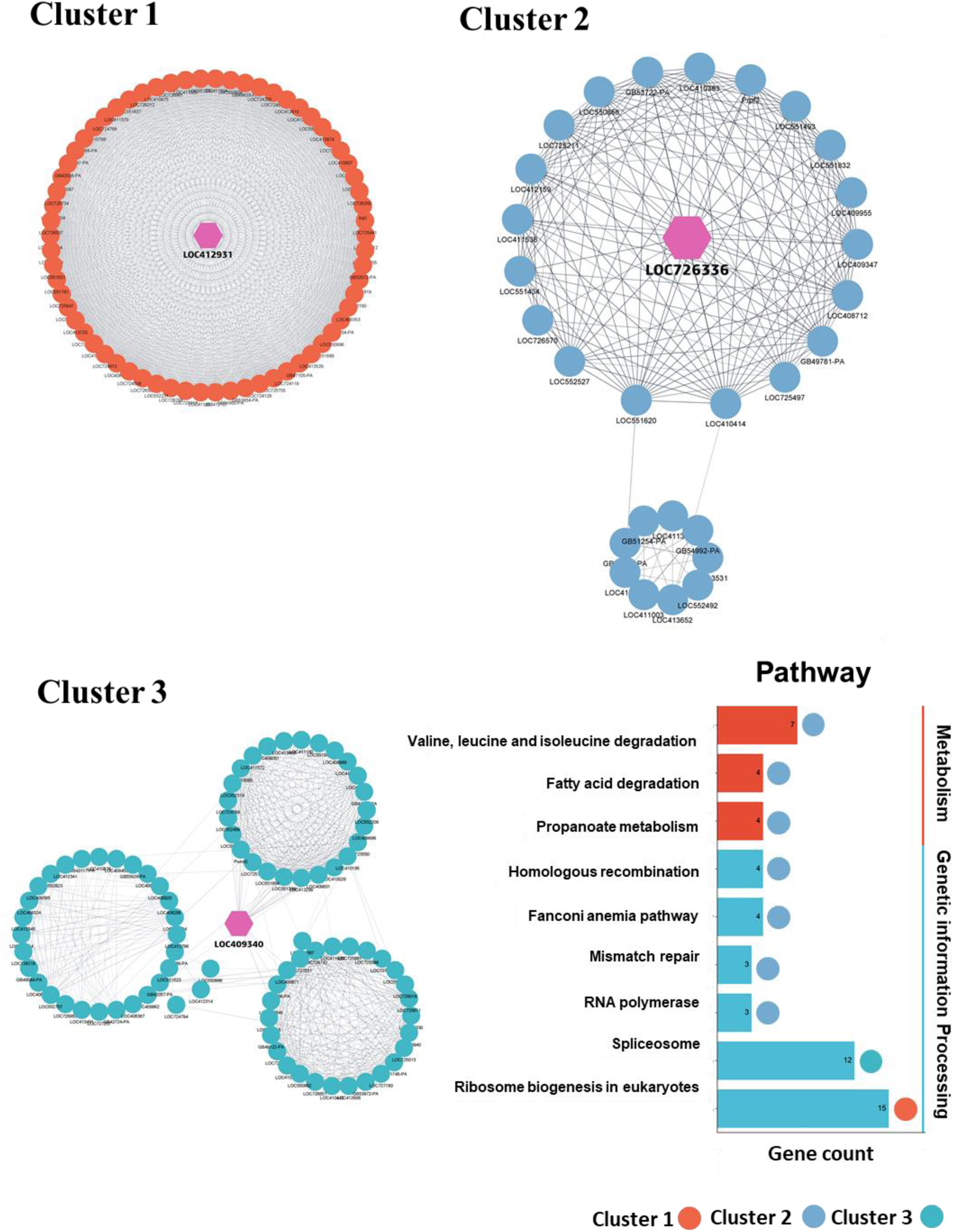
The MCODE clusters were identified from the cDEGs in hypopharyngeal gland tissue under different temperature conditions. The top three modules were considered and grouped into Cluster 1 (Score 62.54); Cluster 2 (Score 15.15); and Cluster 3 (Score 14.07) respectively. Figure generated using Cytoscape.

